# Deciphering the transcriptomic insight during organogenesis in castor (*Ricinus communis* L.), jatropha (*Jatropha curcas* L.) and sunflower (*Helianthus annuus* L.)

**DOI:** 10.1101/679027

**Authors:** Sai Sudha Puvvala, Tarakeswari Muddanuru, Padmavathi AV Thangella, Kumar Aniel O, Navajeet Chakravartty, Saurabh Gupta, Vineeth Kodengil Vettath, Krishna Mohan Ananta Venkata Sri Katta, Sivarama Prasad Lekkala, Boney Kuriakose, Mulpuri Sujatha, Vijay Bhasker Reddy Lachagari

## Abstract

**Background:** Castor is a non-edible oilseed crop with a multitude of pharmaceutical and industrial uses. Profitable cultivation of the crop is hindered by various factors and one of the approaches for genetic improvement of the crop belonging to a monotypic genus is to exploit biotechnological tools. The major limitation for successful exploitation of biotechnological tools is the *in vitro* recalcitrance of castor tissues. Response of castor tissues to *in vitro* culture is poor which necessitated study on understanding the molecular basis of organogenesis in cultured tissues of castor, through *de novo* transcriptome analysis, by comparing with two other crops (jatropha and sunflower) with good regeneration ability.

**Results:** RNA-seq analysis was carried out with hypocotyl explants from castor, jatropha and cotyledons from sunflower cultured on MS media supplemented with different concentrations of hormones. Genes that showed strong differential expression analysis during dedifferentiation and organogenic differentiation stages of callus included components of auxin and cytokinin signaling, secondary metabolite synthesis, genes encoding transcription factors, receptor kinases and protein kinases. In castor, many genes involved in auxin biosynthesis and homeostasis like WAT1 (Wall associated thinness), vacuolar transporter genes, transcription factors like short root like protein were down-regulated while genes like DELLA were upregulated accounting for regeneration recalcitrance. Validation of 62 differentially expressed genes through qRT-PCR showed a consensus of 77.4% with the RNA-Seq analysis.

**Conclusion:** This study provides information on the set of genes involved in the process of organogenesis in three oilseed crops which forms a basis for understanding and improving the efficiency of plant regeneration and genetic transformation in castor.

## Background

Castor (*Ricinus communis* L.) is a tropical plant that belongs to Euphorbiaceae family and grown mainly for its non-edible oil. Despite the premier position India holds with 85% of world’s total castor production dominating international castor oil trade, profitable cultivation of this crop is hampered by the vulnerability of the released cultivars to several biotic threats at various stages of crop growth and the presence of the toxic protein, ricin in the seeds limiting the use of seed cake as cattle feed. The genetic variability to biotic stresses and seed quality traits is limited in the cultivar germplasm [1, 2]. Conventional breeding techniques have limited scope in improvement of resistance to biotic stresses and oil quality necessitating the exploitation of biotechnological and genetic engineering tools [3, 4, 5]. The main prerequisites for genetic improvement are reliable and reproducible protocols of plant regeneration from cultured tissues and a highly efficient transformation system [4, 6].

The morphological, physiological and molecular aspects of *in vitro* shoot organogenesis were studied extensively in various crops. *In vitro* shoot organogenesis is a complex and well-coordinated developmental process which involves several key genes, molecular markers and pathways [7]. In castor, several difficulties were highlighted regarding very slow response for shoot proliferation from the selected organogenic callus cultures [8]. There are only a few reports of plantlet differentiation in castor and in most of the cases, regenerated plantlets were obtained from apical meristems and shoot tip callus [4, 6, 8, 9, 10, 11, 12, 13, 14] and a single report on somatic organogenesis through callus-mediated shoot regeneration [15].

Most of the molecular studies concerned to *in vitro* organogenesis were confined to model plants like Arabidopsis. Che et al. [16] reported hundreds of up and down-regulated genes during *in vitro* callus, shoot and root development in Arabidopsis tissue culture. It is generally thought that pre-incubation on callus induction media is required to permit the dedifferentiation of tissues that will ultimately re-differentiate into organs [17, 18]. Christianson [19] classified the phenomenon of shoot organogenesis into three phases: acquisition of competency, identity specification and differentiation. Valvekens et al. [20] studied indirect organogenesis procedure in Arabidopsis in which root explants were induced to form callus on a callus induction medium and then transferred to shoot induction medium to induce shoots wherein cells become competent to shoot induction signals on callus induction medium itself [21, 22]. There is no single report for gene expression analysis (or) proteomic analysis identifying genes responsible for regeneration recalcitrance in castor. However, tissue-specific whole transcriptome sequencing in castor to understand triacylglycerol lipid biosynthetic pathway to increase ricinoleic acid was reported by Brown et al. [23]. Similarly, not much reports digged into the *in vitro* organogenesis gene pathways in jatropha and sunflower. Studies pertinent to jatropha include transcriptome analysis of flower sex differentiation, reported by Xu et al. [24] wherein the auxin signaling pathway that includes some of the genes like auxin responsive factors, gibberellin-regulated protein, AMP-activated protein kinase to play a major role in stamen development and embryo sac development were identified. Global expression patterns of transcripts regulated by cytokinins in the inflorescence meristems were reported in jatropha [25] and castor [26].

Despite the research efforts that expanded over the past three and half decades in castor tissue culture, no facile protocol of regeneration has been developed so far. Hence, there is an immediate need to understand the molecular basis of *in vitro* recalcitrance in castor. For this, RNA-seq analysis was undertaken as a platform to understand gene expression profiles by comparing the transcript profiles of cultured castor tissues with jatropha (*Jatropha curcas* L.) which is also a member of Euphorbiaceae that shows good regeneration ability [27] and sunflower (*Helianthus annuus* L.), yet another oilseed crop possessing high adventitious shoot regeneration potential [28]. Transcriptome analysis can be a reliable and effective method, which can be used to know the complete set of transcripts in a cell and their quantity, for a specific developmental stage or physiological condition. RNA-seq provides a far more precise measurement of the levels of transcripts and their isoforms than other methods [29]. Furthermore, unraveling these regulatory cascades in castor from the stage of callus induction to shoot regeneration in the plant hormone media would be a major achievement to improve regeneration protocols in castor. Hence, the present study has been undertaken to identify the key genes controlling callus differentiation in castor, understand the molecular mechanism of regeneration in castor by comparing the transcript profiles with other oilseed crops (sunflower, jatropha) proven to have good regenerability *in vitro* as a prelude to overcome the problem of *in vitro* recalcitrance that limits the exploitation of castor through *in vitro* genetic transformation systems.

## Results

### Plant callus generation and sample preparation

In castor, callus-based regeneration was assessed on different combinations and concentrations of growth regulators incorporated in MS media and from explants like roots, cotyledons, hypocotyls, cotyledonary leaves and primary leaves. Of all the media tried, shoot-like structures were observed when cotyledonary leaves from seedlings were inoculated on MS media supplemented with 0.5 mg/l BA+1.0 mg/l NAA. In medium supplemented with 0.5 mg/l TDZ+1.0 mg/l 2-iP+1.0 mg/l NAA, green nodular callus was observed from cotyledon explants. Organogenic callus was observed from hypocotyls on medium supplemented with 0.2 mg/l 2,4-D+2.0 mg/l KN. Brownish green callus was observed from roots on medium supplemented with 0.2 mg/l BA+0.2 mg/l 2,4-D. Green soft callus was observed from the leaf explants when inoculated on media supplemented with 1.0 mg/l TDZ+0.5 mg/l NAA. Brown nodular callus was observed when cotyledons were inoculated on medium with 5.0 mg/l TDZ+2.0 mg/l 2-iP+1.0 mg/l IBA. Green nodular callus was observed from cotyledons inoculated on medium with 2.0 mg/l TDZ+0.1 mg/l IBA. However, good response in terms of green nodular and organogenic callus was observed on medium supplemented with 2.0 mg/l 2-iP+0.1 mg/l TDZ+0.5 mg/l IAA from the hypocotyl explants in castor and jatropha and from the cotyledon explants in sunflower. The response of castor explants on different combinations of growth hormones is illustrated in Figure 1.

**Fig. 1.**
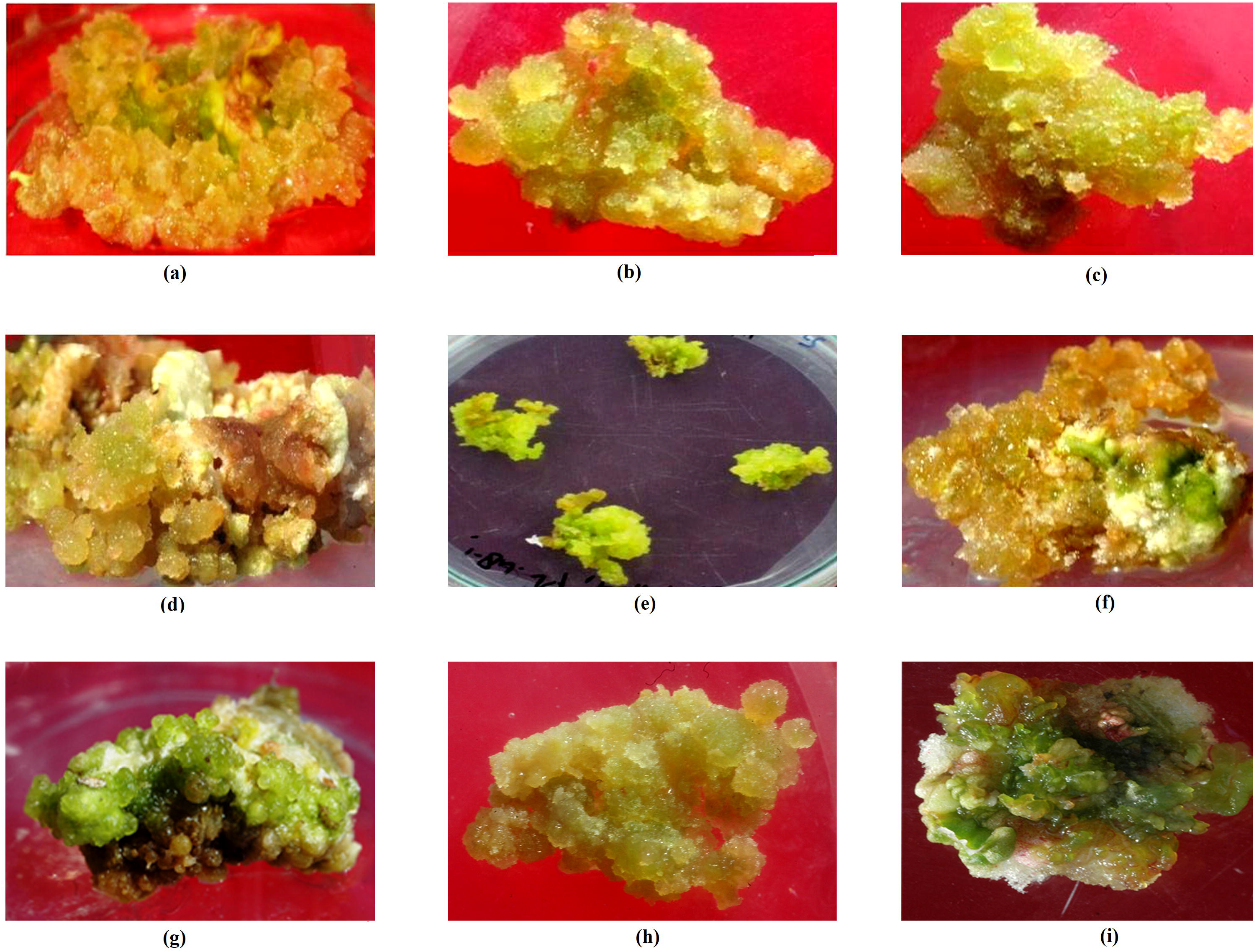
Callus types observed in castor tissues cultured on medium with different combinations of growth regulators. a) Shoot-like structures were observed when leaves were inoculated on medium with 0.5 mg/l BA+1.0 mg/l NAA, b) nodular green callus from cotyledon explants on medium with 0.5 mg/l TDZ+1.0 mg/l 2-iP+1.0 mg/l NAA, c) organogenic callus from hypocotyls on medium supplemented with 0.2 mg/l 2,4-D+2.0 mg/l KN, d) brownish green callus from roots on medium with 0.2 mg/l BA+0.2 mg/l 2,4-D, e) green soft callus from the leaf explants on media supplemented with 1.0 mg/l TDZ+0.5 mg/l NAA, f) brown nodular callus from cotyledons inoculated on medium with 5.0 mg/l TDZ+2.0 mg/l 2-iP +1.0 mg/l IBA, g) green nodular callus from cotyledons on medium with 2.0 mg/l TDZ+0.1 mg/l IBA, h) Good callus observed from leaf explants inoculated on medium with 0.5 mg/l BA+5.0 mg/l 2,4-D + 0.1 mg/l NAA, i) green nodular and regenerable type of callus observed from hypocotyls on medium with 2.0 mg/l 2-iP+0.1 mg/l TDZ+0.5 mg/l IAA.

For RNA-seq analysis, hypocotyl explants from castor and jatropha and cotyledons from sunflower were cultured on MS media supplemented with 2.0 mg/l 2-iP+0.1 mg/l TDZ+0.5 mg/l IAA as this hormonal combination favoured adventitious shoot regeneration in jatropha and sunflower and organogenic callus in castor. In castor, 7 days after culture, explants showed enlargement, however green nodular callus was observed only after 14 days of culture. Retaining the callus on the media resulted only in callus growth but without regeneration. In jatropha, callus formation was observed from the cut ends 7 days after inoculation, the size of the callus was further increased 14 days after culture and after 21 days, shoot like structures were observed from enlarged callus in the explants. In sunflower, the explants showed significant enlargement after 7 days of inoculation, base callus formation with small shoots was observed 14 days after culture and after 21 days, the size of the callus was increased with formation of shoots and their differentiation (Figure 2). Therefore, the samples were collected at time intervals of 0, 7, 14 and 21 days after culture (DAC) for RNA-seq analysis.

**Fig. 2.**
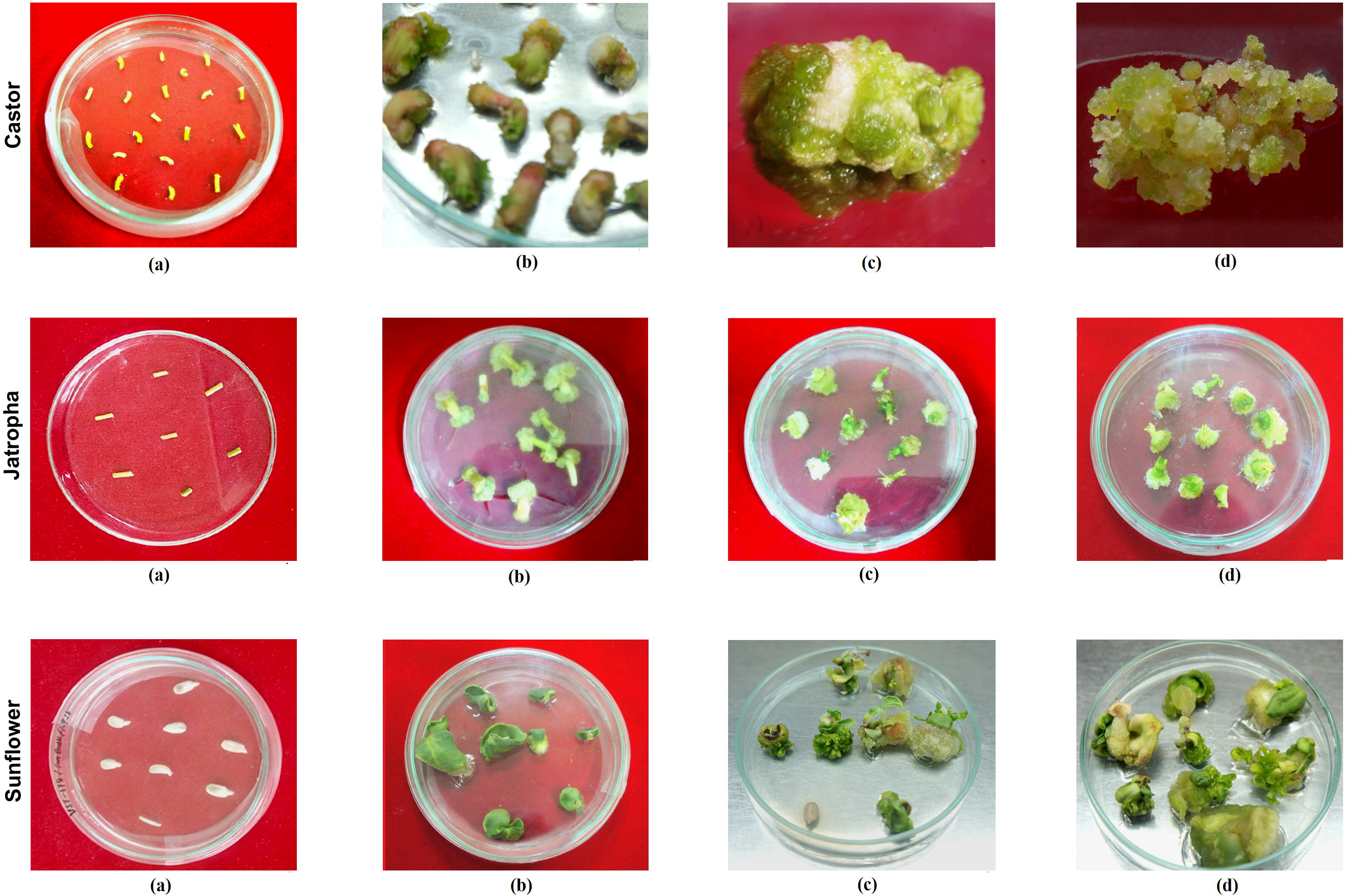
Response of hypocotyls of castor and jatropha and cotyledons of sunflower cultured on medium supplemented with 2.0 mg/l 2-iP+0.1 mg/l TDZ+0.5 mg/l IAA at various time intervals. a, b, c, d represent 0, 7, 14 and 21 days of culture for each of the crops.

### Reads filtering and *de novo* assembly

The quality and quantity of RNA extracted from the control and cultured samples of the three crops were good and cDNA libraries were prepared. A total of 58,304,422 and 44,045,004 raw reads were generated for castor cultured tissues (C-SD) and control castor (CC) samples. The number of raw reads was 5,201,052 and 43,614,842 for jatropha cultured tissues (J-SD) and jatropha control (JC), respectively while 46,548,738 and 38,366,150 raw reads were found in sunflower cultured tissues (S-SD) and sunflower control (SC), respectively (Table 1). The quality of the sequences obtained from the sequencer depends on the base quality score distribution, average base content per read and GC distribution in the reads. The average base quality was above Q30 (error-probability >=0.001) for 92.5% of the bases and GC (%) ranged from 42 to 52 with the highest recorded in the sunflower cultured tissue (Table 1). Along with organellar reads and rRNA, tRNA, snRNA and other RNA reads from the data was also removed from all samples. 32,379,620, 26,099,152, 23,355,268, 30,395,124 and 34,094,516, 35,661,324 filtered reads were obtained in C-SD, CC, J-SD, JC and S-SD, SC, respectively and having GC (%) ranging 44.3 to 52.0, with Q30 range of 95.26 to 96.47 of all filtered samples. High-quality reads were assembled and total of 109,343, 94,073 and 1,30,548 transcripts were annotated in castor, jatropha and sunflower, respectively. Length of longest and mean GC (%) of transcripts (>=200 bp length) of each cultivar (Table 1). Subsequently, 80,092, 62,719, 85,267 unigenes between the control and cultured samples of castor, sunflower and jatropha, respectively were pooled out and further used as transcriptome.

**Table 1.**
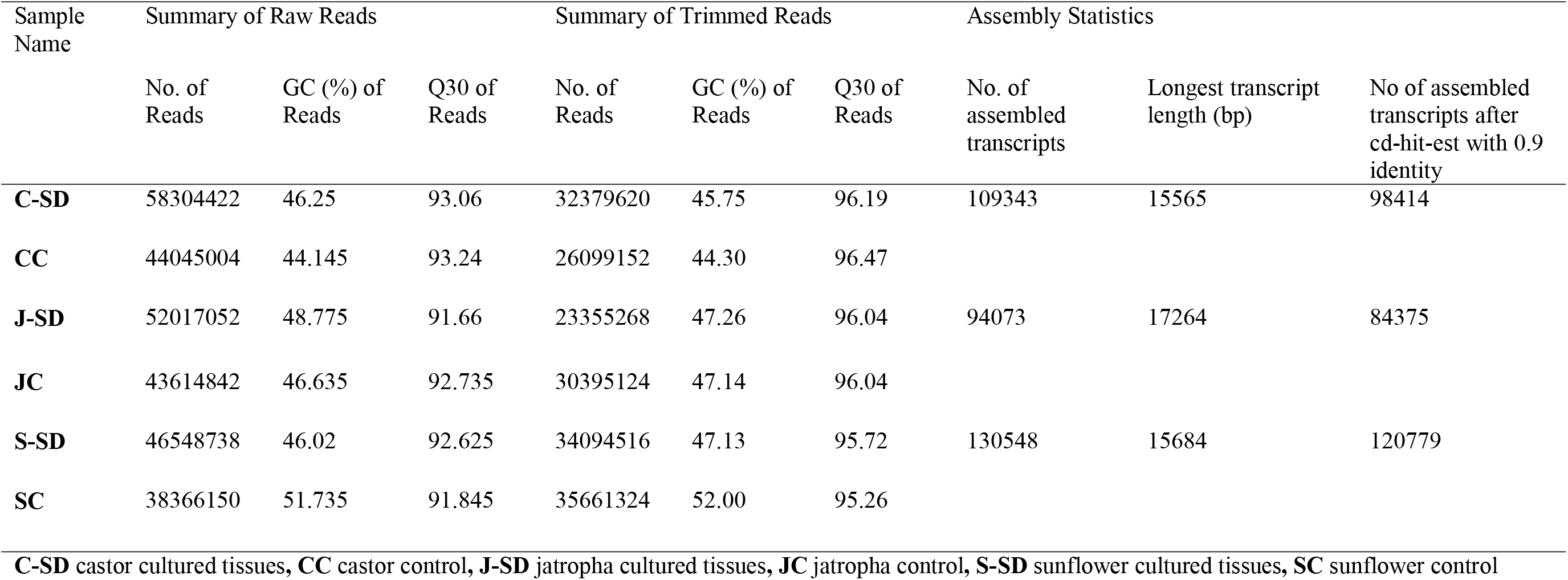
Summary of raw and trimmed reads with assembled reads statistics.

### Estimation of differentially expressed genes

To obtain differentially expressed genes/transcripts (DEGs), trimmed reads of C-SD, CC, J-SD, JC, and S-SD, SC were aligned with assembled transcriptomes of castor, jatropha and sunflower, respectively. Alignment percentages of the reads ranged from 73.6% to 94.2% with the highest alignment percentage observed in castor control tissue (94.2%). A total of 15,194,137 reads were aligned from 161,898,810 filtered paired-end reads in cultured castor samples while 12,297,920 reads were aligned from 13,049,576 filtered paired end reads in the controls (Table 2). Alignment results indicated 98,414, 84,375 and 120,779 unique transcripts (after removal of redundant transcripts) in castor (Additional file 1: Table S1), jatropha (Additional file 2: Table S2) and sunflower, (Additional file 3: Table S3) respectively along with their basic, structural, functional information that has been predicted using BLASTX. It has been identified that 55,576 (69.39%), 40,402 (64.41%) and 52,638 (61.73%) transcripts of castor, jatropha and sunflower have at least one significant hit in NCBI database with identity of 40% at protein level and E-value of >=1e-5 (Table 2). The expression levels were calculated using a normalizing statistic called fragments mapped per kilobase of exon per million reads mapped (FPKM) which provides a measure of expression level that accounts for variation in gene length. A total of 72,416, 57,742, 50,582, 53,627, and 27,416, 75,509 transcripts of C-SD, CC, J-SD, JC, and S-SD, SC, respectively qualified the FPKM ≥1.0 criteria. All unique transcripts were used in EdgeR for DEGs analysis. The bar chart represents the log2fold change (FC) values for all genes, and the cultured samples were compared with control samples (Figure 3). Differential expression analysis of these transcripts based on P values ≤ 0.05 showed 4757, 2325, 738 upregulated and 2630, 1228, 841 downregulated genes in castor (Additional file 4: Tables S4 and S5), sunflower (Additional file 4: Tables S6 and S7) and jatropha (Additional file 4: Tables S8 and S9), respectively. In addition, count per million (CPM) based distribution of DEGs are presented through a volcano plot (Figure 4). DEGs analysis infers that in castor, maximum number of genes is expressed corroborating with our main objective of the study.

**Table 2.** Statistics of read alignment and differentially expressed gene summary.

| Sample identity | No. of filtered reads (paired-end) | No. of reads aligned | Alignment percentage | No. of transcripts$\geq 1$.0 FPKM | Upregulated DGGs | Downregulated DGGs | No. of transcripts with significant BLASTX |
| --- | --- | --- | --- | --- | --- | --- | --- |
| **CC** | 13049576 | 12297920 | 94.24 | 72416 | 4758 | 2631 | 55576 |
| **C-SD** | 16189810 | 15194137 | 93.85 | 57742 |  |  |  |
| **JC** | 15197562 | 14196043 | 93.41 | 50582 | 2326 | 1229 | 40402 |
| **J-SD** | 11677634 | 10383752 | 88.92 | 53627 |  |  |  |
| **SC** | 17830662 | 13135849 | 73.67 | 27416 | 749 | 842 | 52639 |
| **S-SD** | 17047258 | 13883287 | 81.44 | 75509 |  |  |  |

\***C-SD** castor cultured tissues, **CC** castor control, **J-SD** jatropha cultured tissues, **JC** jatropha control, **S-SD** sunflower cultured tissues, **SC** sunflower control

**Fig. 3.**
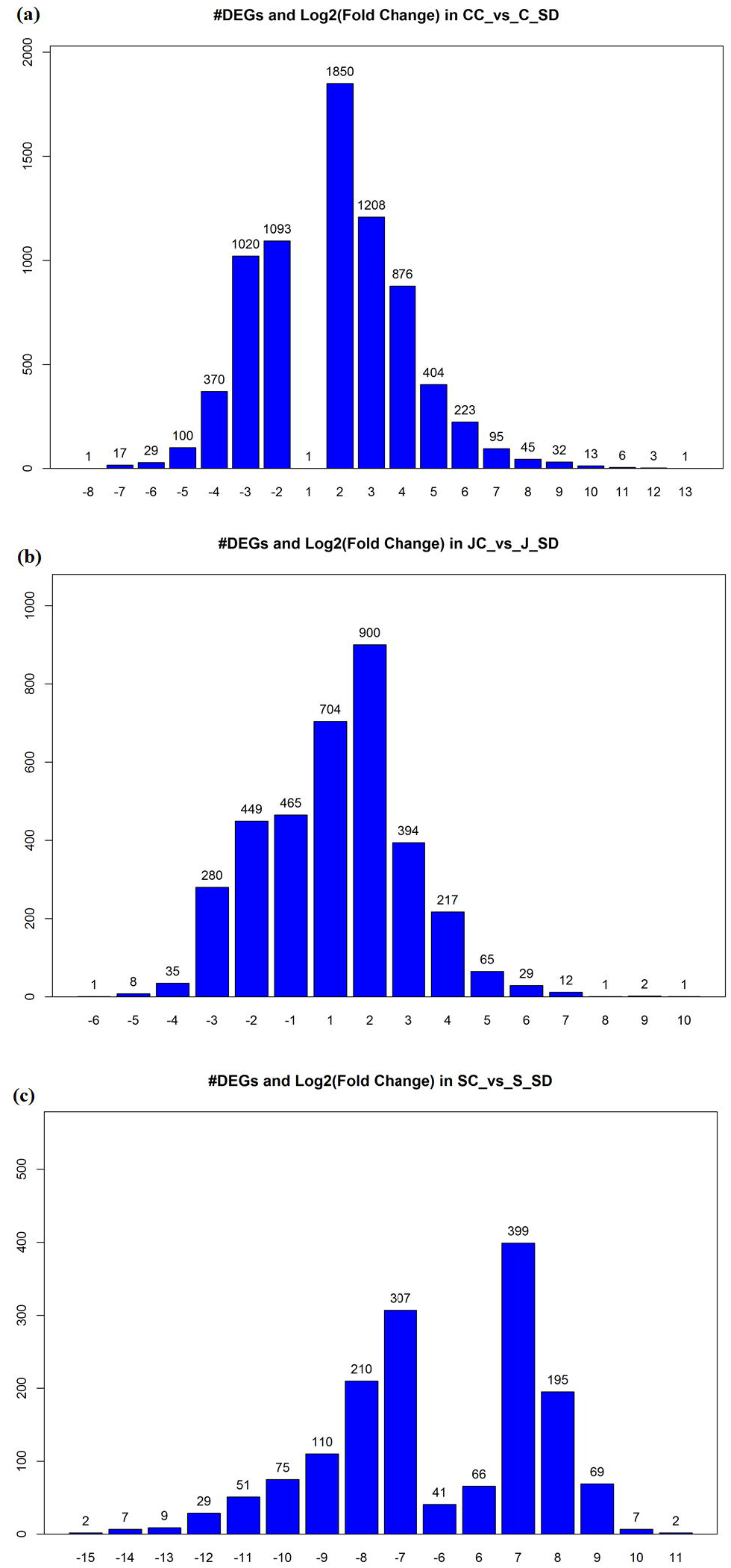
Differentially expressed genes (DEGs) categorized according to log2 Fold Change (FC) values for (a) castor, (b) jatropha, and (c) sunflower.

**Fig. 4.**
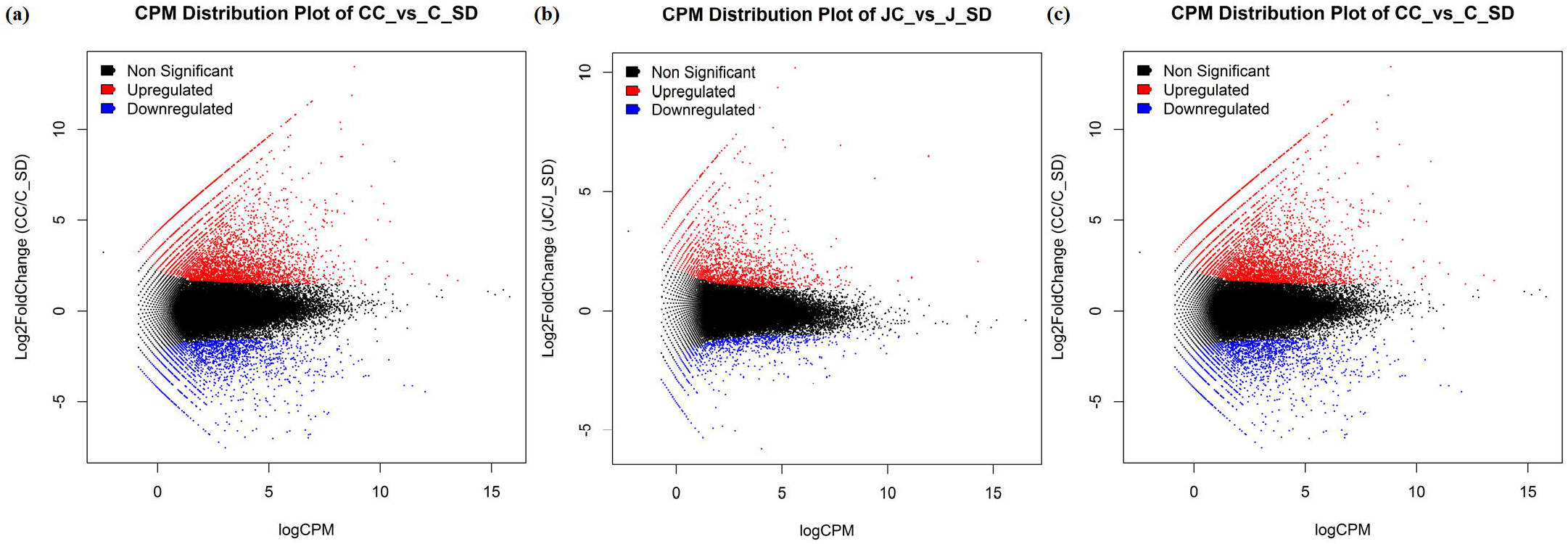
MA plots of differentially expressed genes (DEGs) categorized according to count per million (CPM) values for (a) castor, (b) jatropha, and (c) sunflower.

### Gene ontology and functional annotations of DEGs

The gene ontology (GO) terms for DEGs provides the information about biological processes (BP), molecular function (MF), and cellular components (CC). GO terms for DEGs of castor, jatropha and sunflower with their related information are presented in additional file 4: Tables S4-S9. Moreover, the significant GO terms related to BP, MF and CC in each cultivar are represented in Figure 5. Transcription and regulation of DNA-templated [GO:0006351, GO:0006355], translation [GO:0006412], nucleic acid binding [GO:0003676], carbohydrate metabolic process [GO:0005975], transmembrane transport [GO:0055085], signal transduction [GO:0007165] and protein folding [GO:0006457] terms of BP were prominent in the three crops (Figure 6a) indicating their primary role in regulation of various genes. However, terms carbohydrate metabolic process [GO:0005975], signal transduction [GO:0007165], microtubule-based movement [GO:0007018] and ubiquitin-dependent protein catabolic process [GO:0006511] associated genes are highly expressed in castor in comparison to jatropha and sunflower. MFs such as ATP binding [GO:0005524], zinc ion binding [GO:0008270], integral component of membrane [GO:0016021], nucleic acid binding [GO:0003676], metal ion binding [GO:0046872], protein serine/threonine kinase activity [GO:0004674] terms are the majorly participating in all three crops (Figure 6b). In castor, higher number of genes in DNA binding [GO:0003677], hydrolase activity [GO:0016787], protein kinase activity [GO:0004672], nucleotide binding [GO:0000166], transcription factor activity, sequence-specific DNA binding [GO:0003700], structural constituent of ribosome [GO:0003735], non-membrane spanning protein tyrosine kinase activity [GO:0004715], kinase activity [GO:0016301], iron ion binding [GO:0005506], sequence-specific DNA binding [GO:0043565], ligase activity [GO:0016874] and calcium ion binding [GO:0005509] terms are associated in comparison to jatropha and sunflower. Similarly, the CCs terms such as integral component of membrane [GO:0016021], ATP binding [GO:0005524], DNA binding [GO:0003677], intracellular [GO:0005622], nucleus [GO:0005634], ribosome [GO:0005840], cytoplasm [GO:0005737] terms associated genes are highly expressed in the three crops (Figure 6c). Overall this analysis provides a basic idea of gene function and further, in-depth analysis of gene function was carried out for DEGs and top hits were considered to extract organism name as well as their functions. In castor, *Ricinus communis* L. occupied the first place with highest number of transcripts followed by *Jatropha curcas, Vitis vinifera, Populus trichocarpa*, and *Populus euphratica*. In case of Jatropha, the number of transcripts were highest in *Jatropha curcas*, followed by *Ricinus communis* and *Vitis vinifera*. *Cynara cardunculus* var. *scolymus* had the maximum hits followed by *Vitis vinifera*, *Sesamum indicum*, and *Helianthus annuus* in case of sunflower. Functions of DEGs were annotated using UniProt database and listed in Additional file 4: Tables S4-S9 with their functional and structural descriptions.

**Fig. 5.**
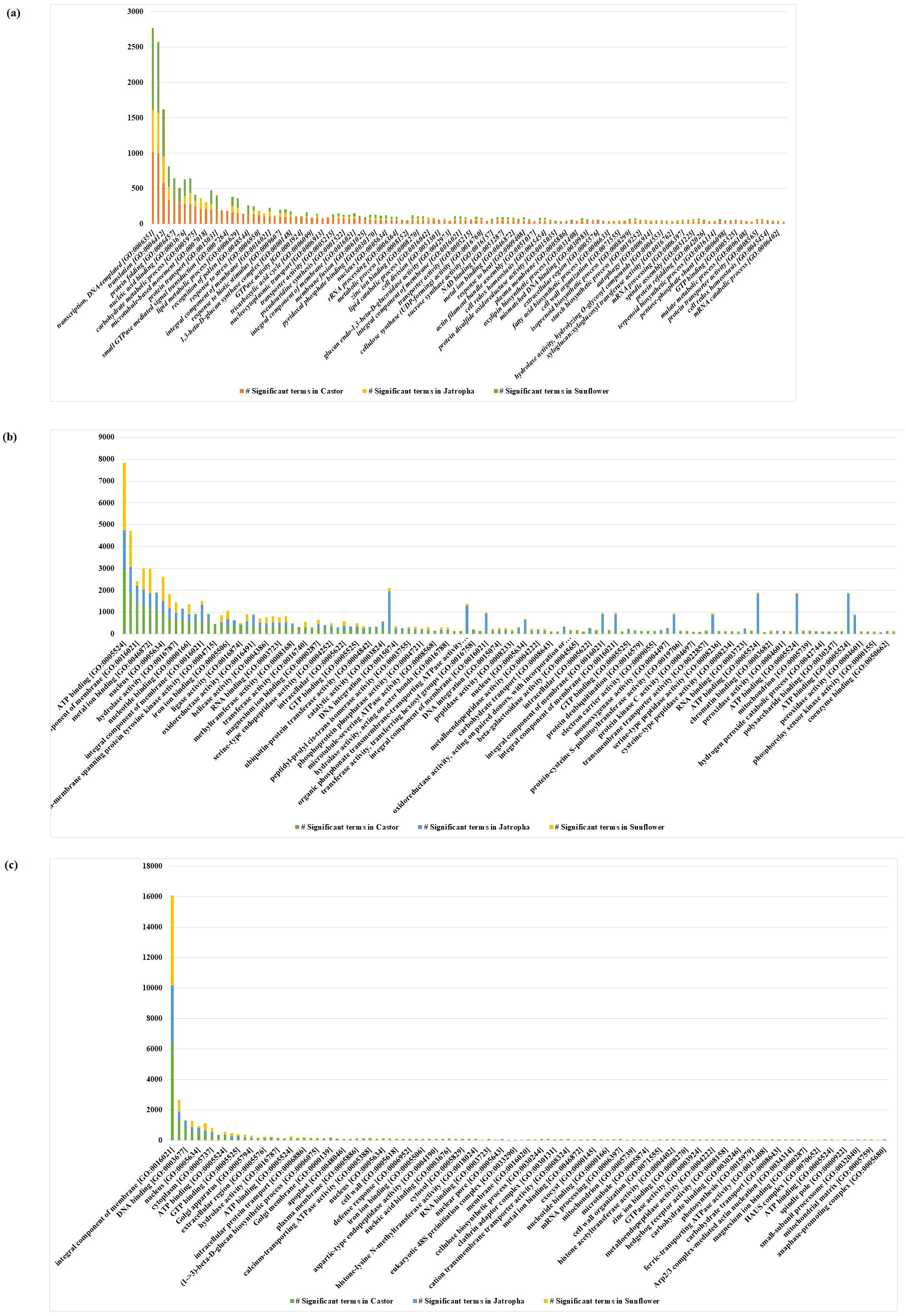
Significant enriched Gene Ontology (GO) terms associated with different (a) biological processes, (b) molecular functions, and (c) cellular components. GO terms have been represented for castor, jatropha and sunflower in different colours below each category of the figure.

**Fig. 6.**
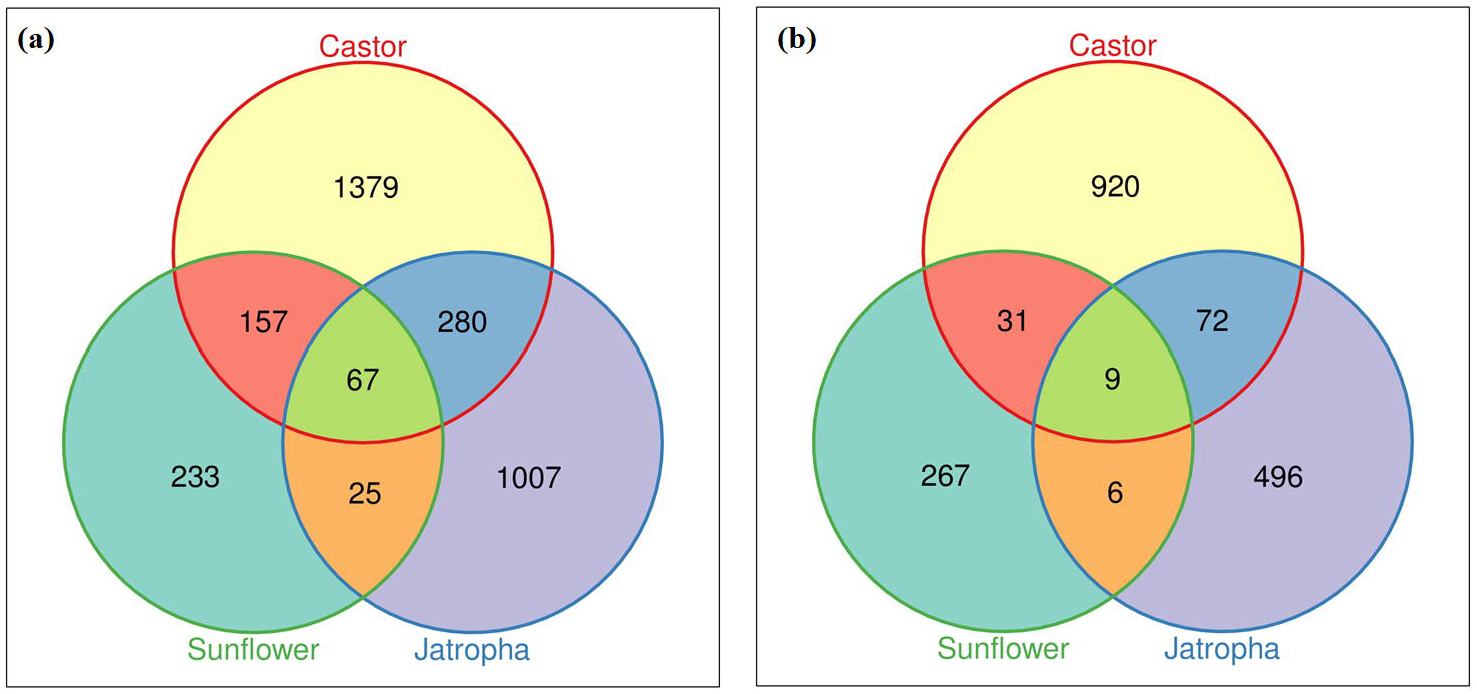
Venn diagram indicates the common and unique orthologues groups identified for (a) upregulated, (b) downregulated DEGs from castor, jatropha and sunflower.

### DEGs involved in callus formation, plant growth and hormone metabolism

Genes that play an important role in auxin biosynthesis were observed in the three crops. Particularly, in castor, higher number of genes that play critical role in maintaining auxin levels were found to be downregulated while comparing with jatropha and sunflower. Auxin-induced protein 15A, indole-3-acetic acid-induced protein, IAA-amino acid hydrolase ILR1-like 3, auxin-induced protein, putative required for IAA biosynthesis, wall associated thinness (WAT1), vacuolar transporter genes, hypothetical proteins were significantly down-regulated. Noticeably, not many cytokinin pathway related genes were expressed in castor except, cytokinin riboside 5’-monophosphate phosphoribohydrolase, histidine-containing phosphotransfer protein which showed down-regulation (Additional file 4: Tables S4 and S5). In jatropha, axial regulator (YABBY 1) gene that regulates the initiation of the embryonic shoot apical meristem (SAM) development, auxin transporter-like protein 3, a carrier protein which is involved in proton-driven auxin influx and mediates the formation of auxin gradient from developing leaves (site of auxin biosynthesis) to tips were up-regulated. Besides, cytokinin riboside 5’-monophosphate phosphoribohydrolase LOG5-like which is very important for shoot regeneration is up-regulated. The gene abscisic acid 8’-hydroxylase1 is found to be upregulated in castor and jatropha and catalyzes the committed step in the major ABA catabolic pathway (Additional file 4: Tables S6 and S7). However, in sunflower, significant plant hormone biosynthesis genes were not identified (Additional file 4: Tables S8 and S9).

### DEGs involved in different binding and cellular transportation activities

In plants, various metals and biomolecules binding activity is generally supported by cellular transportation to maintain the cellular internal and external integrity. In castor, amino acid binding proteins, ATP binding proteins, metal ion binding proteins, calmodulin binding proteins, chlorophyll A/B binding proteins, DNA binding proteins, dehydration-responsive element-binding proteins, GTP binding proteins, lipid binding proteins and nucleic acid binding proteins were upregulated, while boron transporters, ATP-binding cassette transporters, calcium-transporting ATPases, copper transporters, sugar transporters, protein binding proteins and oligopeptide transporters also showed upregulations. Besides, downregulated binding and transporter proteins in castor include calcium ion binding protein, calmodulin binding protein, metal ion binding protein and ABC transporter family proteins, aquaporin transporters, ATP-binding cassette transporters, transporter proteins, Alanine-glyoxylate aminotransferase proteins, benzoate carboxyl methyltransferase, cationic amino acid transporter, copper-transporting ATPase, bidirectional sugar transporter, oligopeptide transporter, glycosyltransferase. The higher expression of ABC transporter family proteins binding and transporter, ATP-binding cassette transporters copper and sugar transporters indicates their role in growth regulator transportation (Additional file 4: Tables S4 and S5) [30, 31]. Similarly, various binding and transportation activities also observed in jatropha and sunflower and related protein expression can be seen in additional file 4: Tables S6, S7 and Tables S8 and S9, respectively.

### Identification of DEGs that work as transcription factors and other important proteins

Transcription factors are the regulatory proteins which upon binding to specific DNA sequences regulate target gene expression. In castor, some of the transcription factors up-regulated belong to WRKY TF while GATA TF, NAC domain containing protein 62, zinc finger proteins, r2-r3 myb TFs, protein short root like were down-regulated. In jatropha, up-regulated TFs include MYB family (MYB 108, MYB 308), homeobox-leucine zipper protein MERISTEM L1 involved in the cell specification and pattern formation during embryogenesis, WRKY family (WRKY transcription factor 75 isoform X1). However, wound-induced protein, WIN1 precursor that favors callus formation from wound tissue was down-regulated. Further, some important proteins participating in general plant stress defense mechanism such as, Heat shock proteins family, Flavonol synthase/flavanone 3-hydroxylase, fructose-bisphosphate aldolase, disease resistance response proteins, etc. were also upregulated in castor [32, 33] while serine proteinase family proteins are found downregulated in castor [34]. Along with many other hypothetical proteins may also participate in plant stress defense mechanism [35].

### Identification of DEGs involved in signal transduction

In jatropha, many of the genes involved in signal transduction were significantly up-regulated. These involve many probable LRR receptor-like serine/threonine-protein kinases, receptor-like protein kinase, receptor-like protein 12, serine/threonine-protein kinases SAPK1, Calcium-dependent protein kinase [36, 37]. This clearly implies the signaling cascade events downstream the genes involved in organogenesis. In contrary, the brassinosteroid LRR receptor like kinases were down-regulated in castor.

### Cell wall related genes

In castor, cell wall related genes that are up-regulated include pectinesterase-2 precursor involved in the dimethylesterification of cell wall pectin, polygalacturonase non-catalytic subunit AroGP2 precursor involved in cell wall organization, beta expansin 3 in loosening of cell walls poor in pectin and xyloglucans, o-methyltransferase in monolignol biosynthesis. Down-regulated genes are non-specific lipid-transfer protein 1-like proteins which are small, basic proteins that have been reported to be involved in numerous biological processes such as transfer of phospholipids, and reproductive development. Another gene, lupeous synthase responsible for formation of the cuticular lupeol conferring characteristic surface properties of *R. communis* stems is also down-regulated.

### Genes involved in biosynthetic pathways

In castor, some of the up-regulated genes were found to play a role in secondary metabolite biosynthesis pathways like reticuline oxidase in biosynthesis of isoquinoline alkaloid biosynthesis, cycloartenol synthase in sterol biosynthesis, muconate cycloisomerase in benzoate degradation. Flavonol synthase/flavanone 3-hydroxylase involved in flavonoid biosynthesis, sesquiterpene synthase, (R)-limonene synthase involved in monoterpenoid biosynthesis were down-regulated. Cytochrome P450 which is up-regulated 10-fold is involved in ursolate biosynthesis. Ursolate is a pentacyclic triterpenoic acid that occurs naturally in many plants. Thaumatin-like proteins are related to highly complex gene family involved in a wide range of developmental processes in fungi, plants, and animals.

### Identification of orthologous genes group from DEGs

In the orthologous gene analysis, a total of 67 gene groups were commonly up-regulated in all the three crops (Additional file 4: Tables S10-S12), while 1,379 genes groups were uniquely expressed in castor, 1,007 genes group in jatropha and 233 genes group in sunflower. Both castor and sunflower share 157 up-regulated genes group in common and absent in jatropha, while 280 genes group were commonly expressed only between castor and jatropha and absent in sunflower. Only 25 genes group were commonly expressed between sunflower and jatropha (Figure 6). A total of nine groups were commonly down-regulated in all the three crops (Additional file 4: Tables S13-S15) while 920 genes group were uniquely down-regulated in castor, 496 genes group in jatropha and 267 genes group in sunflower. Thirty-one genes group were expressed in both castor and sunflower and absent in jatropha, while 72 genes group were commonly expressed only between castor and jatropha and absent in sunflower. Only 6 genes group were commonly expressed between sunflower and jatropha. The significantly down-regulated genes group in castor include phenylpropanoid pathway genes group like jasmonate O-methyltransferase, cinnamoyl-CoA reductase, O-methyltransferase, receptor kinases like Brassinosteroid Insensitive 1-associated receptor kinase 1, homeobox protein knotted-1, hormone biosynthesis genes like somatic embryogenesis receptor kinase, auxin-responsive protein, auxin efflux carrier component, indole-3-acetic acid-amido synthetase GH3.1, GRAS13 protein, gibberellin-regulated protein 1, gibberellin receptor GID1, oilseed pathway genes like delta 9 desaturase, transcription factors like WRKY transcription factor 16, Transcription factor TGA7, 9-cis-epoxycarotenoid dioxygenase and cell wall related genes like glycine-rich cell wall structural protein 1, WAT1-related protein, etc. The significant up-regulated genes include transcription factors like GATA transcription factor, R2r3-myb transcription factor, WRKY transcription factor, CBF-like transcription factor, ethylene-responsive transcription factor, hormone biosynthesis genes like stem 28 kDa glycoprotein, gibberellin receptor GID1, chitin-inducible gibberellin-responsive protein, SAUR-like auxin-responsive protein, auxin:hydrogen symporter, stem-specific protein tsjt1, DELLA protein GAIP-B, phenazine biosynthesis protein, transporter genes like UDP-sugar transporter, potassium channel KAT3. Overall this analysis provided commonly upregulated and downregulated genes for *in-vitro* validation.

### qRT-PCR validation of DEGs

Seventy-two DEGs by selecting some of the most significantly upregulated and downregulated genes were subjected to qRT-PCR analysis. Of the tested genes, primers for 10 genes failed to produce amplification. Of the remaining 52 genes, qRT-PCR analysis of 48 genes was in close agreement with the RNA-Seq data. Figure 7 represents data from two each of the upregulated and downregulated genes of castor (a-d), Jatropha (e-h), sunflower (i-l) and other genes reported to have known function in organogenesis (m-t). Amplification was observed in all the three crops, two crops or a single crop. Perusal of the data presented in Figure 7 and additional file 5: Table S16 show that the differential expression in terms of upregulation or downregulation is consistent at all time points, (eg: Fig 7a, b, d for castor); upregulated followed by downregulation (Fig. 7c, g, k for castor, jatropha and sunflower, respectively) and *vice versa* (Fig. 7j, k, i for castor, jatropha and sunflower, respectively).

**Fig. 7.**
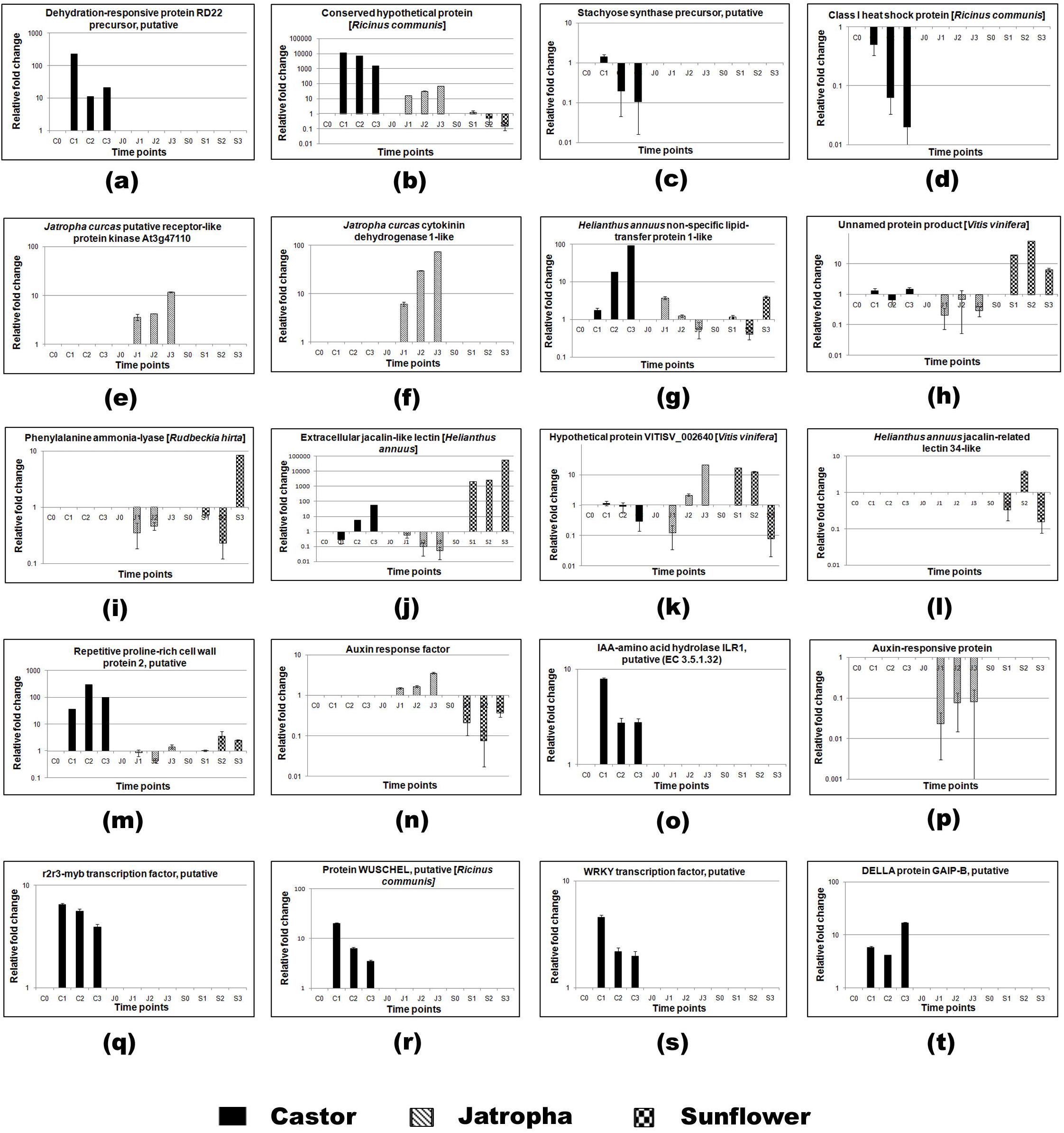
Relative fold change of upregulated and downregulated genes through quantitative real time PCR in the tissues of castor (C0-0 day, C1-7 days, C2-14 days, C3-21 days), jatropha (J0-0 day, J1-7 days, J2-14 days, J3-21 days) and sunflower (S0-0 day, S1-7 days, S2-14 days, S3-21 days) after inoculation. **a, b** represent upregulated genes of castor, **c, d** represent downregulated genes of castor, **e, f** represent upregulated genes of Jatropha, **g, h** represent downregulated genes of Jatropha, **i, j** represent upregulated genes of sunflower, **k, l** represent downregulated genes of sunflower, **m-t** represent other genes reported to govern organogenesis.

## Discussion

Callus is an unorganized, undifferentiated mass of cells with root and shoot primordials produced from a single differentiated cell and many callus cells exhibit totipotency [38, 39]. Appropriate combination of plant growth hormones in tissue culture media makes plant cells exhibit properties like cellular totipotency, developmental plasticity and subsequent regeneration into mature plants [40, 41, 42]. The ratio of auxins-to-cytokinin in the growth media primarily decides the developmental fate of a regenerating tissue *in vitro.* Usually, a higher auxin-to-cytokinin ratio favors root formation, an intermediate ratio promotes callus induction, while a high ratio of cytokinin-to-auxin promotes shoot regeneration [40]. There are two modes of regeneration for a plant cell *in vitro*; a) direct/meristem based and indirect/callus based [43]. A direct organogenesis mode does not require a de-differentiation phase wherein the explants are fully competent while callus-based regeneration involves two steps as described by Motte et al. [44]. Callus induction requires the interplay of several key regulators of auxin and cytokinins signaling pathways and spatially and temporally controlled intrinsic developmental programmes interconnected at multiple levels. Auxins mediate founder cell specification, the development of primordia and the acquisition of organogenesis competence while cytokinins assign shoot identity to the developing primordia.

Plant hormones act as first messengers in regulating the activity of gene *via* various signaling pathways and initiate gene expression. Although various media experimented in our tissue culture studies induced callus from cultured explants of castor with a low frequency, shoot differentiation was sporadic with low reproducibility which implies that the calli contain very few morphogenic cells interspersed in several non-morphogenic tissues. Sujatha and Reddy [45] assessed the morphogenic competence of castor tissues on several basal media supplemented with many growth regulators individually and in a broad range combination according to De Fossard et al. [46] with 81 combinations of minerals, sucrose + growth factors + amino acids besides growth regulators, which revealed low caulogenic response of castor explants for direct as well as callus mediated shoot regeneration. In castor, i*n vitro* propagation system based on hypocotyl derived callus cultures was developed [8]. Hence, this poor response to *in vitro* culture necessitated the study on understanding the genes that are transcribed in cultured tissues of castor by comparing with two other crops (jatropha and sunflower) possessing good regeneration ability through *de novo* transcriptome analysis.

Tissue-culture based plant cells have unique ability to reprogrammed differentiated somatic cells to de-differentiate, proliferate and re-differentiate into whole plants [47]. The gene ontology assignments of the three crops indicated that most of genes are concerned with transcription, regulation of transcription and translation indicating steady state gene expression and reprogramming differentiated cells. For plant cells to transit from callus to shoot organogenesis, they need to remodel their gene expression during which several genes associated with cell division, stress response, primary metabolism and cell wall synthesis get involved [48]. This is done by cellular dedifferentiation i.e., acquiring cellular totipotency which makes cells re-enter into the cell cycle making remarkable changes in the pattern of gene expression as cells switch from one somatic cell to a new one directing either reentry into the cell cycle, cell death, or trans- or dedifferentiation.

The calli or very young primordials can respond to signals that direct the formation of an organ [49, 50, 51]. A cross talk between auxins and cytokinins is required for patterning of the shoot primordium and the shoot meristem [52, 53, 54, 55]. Auxin is very important for positioning of root stem cell niche, shoot and root organogenesis [55, 56, 57]. In castor, auxin-induced protein 15A and Indole-3-acetic acid-induced protein ARG7 that belong to a large auxin responsive gene family, SAURs (Small Auxin Up RNA) [58] were down-regulated 5-folds while IAA-amino acid hydrolase ILR1-like 3 was upregulated. These genes are involved in tryptophan dependent IAA biosynthesis. Indole-3-acetic acid (IAA) is the most abundant naturally occurring auxin in plants that is required throughout the development process. These auxin levels are regulated naturally by forming IAA (Indole-3-acetic acid) conjugates with some of the amino acids. IAA is the most abundant naturally occurring auxin in plants which acts in every aspect of plant development. IAA-amino acid hydrolase ILR1-like 3 is an enzyme which cleaves the conjugates and releases free IAA. Enhanced expression of this enzyme in castor might be one of the reasons for making IAA available in castor cells favoring callus and root growth instead of shoot differentiation [42, 59]. Another gene, auxin-induced protein 5NG4, putative, which is highly and specifically induced by auxin in juvenile shoots prior to adventitious root formation exhibited down-regulation indicating insufficient synthesis of IAA necessary for shoot initiation. Further, down-regulation of genes that play a major role in maintaining auxin homeostasis like WAT1 (Wall associated thinness), vacoular transporter genes inhibit the development of cell wall components and hence, shoot and root morphogenesis. After auxins, high cytokinin levels determine the shoot identity of the organ primordia by establishing a shoot stem cell niche [60]. A first prerequisite for shoot formation is that the cytokinins from the shoot induction media reach the cells that have acquired organogenic competence. Histidine-containing phosphotransfer protein is involved in biosynthesis of the cytokinin, zeatin. These cytokinins are responsible for cell division and shoot initiation. It is down-regulated 4-fold in castor. In jatropha, auxin transporter-like protein 3 which is a carrier protein involved in proton-driven auxin influx mediates the formation of auxin gradient from developing leaves (site of auxin biosynthesis) to tips. These auxin influx and efflux carriers maintain local auxin maxima, essential for shoot regeneration [61, 62]. Up-regulation of some of the genes like Xyloglucan endotransglucosylase/hydrolase synthesizing xyloglucan polymers, essential constituent of the primary cell wall, participating in cell wall construction of growing tissues and 14 kDa proline-rich protein DC2.15 initiating embryogenesis might be responsible for higher regeneration potential in jatropha.

In Arabidopsis, Wuschel protein which is a homeobox transcription factor is expressed during embryogenesis and organogenesis leading to the proliferation of mersitematic tissue from vegetative organs. Localized expression of this protein is considered as a reliable marker for shoot regeneration in Arabidopsis [51] and Medicago [63]. However, in the present study, activity of Wuschel protein is lowered 3 and 7-folds (Additional file 5: Table S16) by the 14^th^ and 21^st^ day of culture, respectively implying less regeneration potential. Li et al. [14] correlated WUS expression with the budding rate from castor epicotyls and found that the expression varied with concentration of the cytokinin and the pre-treatment duration. While in the present study, WUS expression was found to drastically decline in the dedifferentiated tissue. NAC proteins are one of the largest groups of plant-specific transcription factors and are known to play essential roles in various developmental processes, auxin signaling and postembryonic shoot meristem formation [64]. Protein short root like is a transcription factor required for quiescent center cells specification and maintenance of surrounding stem cells and for the asymmetric cell division involved in radial pattern formation in roots. It regulates the radial organization of the shoot axial organs and is required for normal shoot gravitropism [65]. Hence, the down-regulation of these transcription factors also could have contributed to recalcitrance in castor. Transcription factors of the Apetala2/Ethylene Response Factor (AP2/ERF) family like wound-induced dedifferentiation 1 (WIND1) were found to trigger cell dedifferentiation and proliferation leading to callus formation [66]. This clearly shows that auxin perception and the activation of several auxin signaling modules simultaneously is required for shoot organogenesis. Hence, the defects in auxin signaling would probably cause regeneration recalcitrance.

Protein kinases are enzymes that catalyze the transfer of phosphate groups from a nucleoside triphosphate to amino acids such as serine and threonine or histidine residues present in plant proteins thereby modulating the properties. The receptor kinase activation is the starting point of the signaling cascade mediating developmental switches/hormone responses; it represents an important regulatory control point. In jatropha, up-regulation of these protein kinases lead to active signaling for shoot organogenesis. Probable LRR receptor-like serine/threonine-protein kinase, together with RPK2 is required for pattern formation along the radial axis i.e., the apical embryonic domain cell types that generate cotyledon primordia and the apical-basal axis. Other significant proteins like EXORDIUM-like 2 protein plays an important role in brassinosteroid-dependent regulation of plant growth and development, Thaumatin-like proteins are related to highly complex gene family involved in a wide range of developmental processes in fungi, plants, and animals, alpha carbonic anhydrase 1, chloroplastic-like carry inorganic carbon for actively photosynthesizing cells [67]. Phospholipase A1-II 1-like proteins are major digestive enzymes and play a critical role in most physiological processes including the generation of numerous signaling lipids.

## Conclusions

Overall this study deals with organogenic differentiation *in vitro* in three oilseed crops; castor, jatropha and sunflower, castor proved to be highly recalcitrant to *in vitro* manipulations despite research over the past three and half decades and showed extremely low percentage of caulogenic ability from the induced callus and successive plant regeneration. Hence, the investigations undertaken to unravel the reason for recalcitrance in castor using transcriptomic analysis revealed the prime reasons to be the imbalance in auxin metabolism, leading to insufficient accumulation of auxins essential for shoot regeneration. Further, transcription factors like Wuschel responsible for promoting shoot regeneration, protein short root like contributing shoot organogenesis and histidine containing phosphotransfer protein involved in cytokinin biosynthesis for promoting shoot regeneration are down-regulated. Strikingly, there were no signaling cascades activated to promote any shoot regeneration as there seems to be down-regulation of Brassinosteroid LRR receptor kinases. Noticeably, many secondary metabolite synthesis genes were up-regulated in castor. While, looking into the expression patterns of jatropha, many kinases involving in signal transduction were up-regulated indicating a possible role in shoot organogenesis processes besides those involved in cellular processes. In addition, auxin and cytokinin biosynthesis genes were also up-regulated. In sunflower, most of the genes expressed belong to those involved in cellular processes, biochemical pathways and photosynthesis. The interplay between type, amount and timing of growth regulators stimulating genes encoding proteins for hormone synthesis, transport and signaling and their positive and negative regulations play a major role in organogenesis response. Hence, the negative regulatory mechanisms in castor and positive regulation in jatropha and sunflower may be attributed for the major differences in organogenic response observed in this investigation. Our data complements further investigations and gene validations on a broader panel of genotypes and tissues cultured on media with different growth regulators for overcoming the problem of recalcitrance *in vitro* in castor.

## Methods

### Plant material and culture conditions

The seeds of the three oilseed crops; castor, sunflower and jatropha were obtained from ICAR-Indian Institute of Oilseeds Research (IIOR), Hyderabad and the varieties used were DCS–107 for castor, DRSH-1 for sunflower and JP-2 for jatropha.

### Tissue culture studies

Decoated seeds from all the three crops were surface sterilized and inoculated on ½ strength Murashige and Skoog [68] media for germination and growth. From these seedlings, explants like root, hypocotyl, cotyledonary leaf and primary leaf were taken, cut into 0.5 cm size and inoculated onto MS agar medium supplemented with different concentrations and combinations of growth regulators [benzyladenine (BA) + naphthaleneacetic acid (NAA); 2,4, dichlorophenoxyacetic acid (2,4-D) + kinetin (KN); BA+2,4-D+NAA; thidiazuron (TDZ) singly or in combination of 2-isopentenyl adenine (2-iP) with auxins NAA or indole-3-butyric acid (IBA) or indole-3-acetic acid (IAA)] for callus induction and shoot regeneration. The inoculated cultures were maintained at 27 ± 1 ^0^C under a 16/8 hr light/dark photoperiod with light intensity of 30 μmol m^−2^ s^−1^. Of the various media combinations tested, good regenerable callus was observed on medium supplemented with 2.0 mg/l 2-iP+0.1 mg/l TDZ+0.5 mg/l IAA from hypocotyl explants in castor and jatropha and cotyledonary explants of sunflower. A common medium was selected for the three crops to minimize the differences in gene expression due to exogenous growth regulators. After culture initiation the callus was collected at 0^th^ day known as control samples (CC, JC, SC) and 7^th^, 14^th^ and 21^st^ days calli were pooled for organogenesis study and denoted as cultured samples (C-SD, J-SD, S-SD) of castor, sunflower and jatropha.

### Total RNA extraction from samples

Three biological replicates of callus tissue and regenerating explants (about 50-100 mg) were collected from control and cultured hypocotyl explants in castor and jatropha and from cotyledonary explants in sunflower. Further, all samples were washed with DEPC water and immediately frozen at −80 °C. The tissue was crushed into fine powder in liquid nitrogen and RNA was isolated as described in Qiagen RNA extraction kit. The quality of RNA was checked on 1% agarose gel and evaluated on Nanodrop ND 1000 spectrophotometer (Genway, USA). After ensuring the quality (1.8-2.0 at A_260_/A_280_nm) and concentration (250-300 ng/µl), total RNA were used for library preparation.

### Library construction and RNA-sequencing

Quality of extracted total RNA was assessed using Agilent 2100 Bioanalyzer. Those having RNA Integrity Number (RIN) value ≥ 8 is considered as good quality. Approximately 4 ug of total RNA was used to prepare the RNA-seq library using the TruSeq RNA Sample Prep Kits (Illumina) as per the kit protocol. In short, poly-A containing mRNA molecules were purified using poly-T oligo-attached magnetic beads. Following purification, the mRNA was fragmented into small pieces using divalent cations under elevated temperature. The cleaved RNA fragments were used to synthesize first strand cDNA using reverse transcriptase and random primers followed by second strand cDNA synthesis using DNA polymerase I and RNase H. These cDNA fragments were subjected to an end repair process with the addition of a single ‘A’ base, and then ligation of the adapters. The products were purified, and PCR enriched to create the final cDNA library. Bioanalyzer plots were used at every step to assess mRNA quality, enrichment success, fragmentation sizes, and final library sizes. Both Qubit and qPCR were used for measuring the quantity of the library before sequencing. After the libraries were constructed, they were sequenced on HiSeq-2500 to obtain 2 x100 bp paired ends having 40 million high quality reads/sample. The parameters used to check the quality depend on the base quality distribution, average base content per read and GC distribution.

### *De novo* transcriptome assembly

Fastq files of all samples (control and cultured) were preprocessed before performing the assembly. The adapter sequences were trimmed, and the reads were also filtered out wherever the average quality score was less than 20 in any of the paired end reads. The high-quality reads were then assembled using Trinity v2.02 [69] with default options. Redundancy of the transcript fragments were minimized using cdhit-est v4.6 [70]. The GC content distribution of all the assembled transcripts was calculated. Transcripts of length >= 200 bp were found ideal and considered for transcript expression estimation and downstream annotations. The transcriptome sequences of control and cultured samples of castor, jatropha and sunflower as raw reads are submitted to NCBI and can be accessed with the NCBI accession number PRJNA415556.

### Gene expression estimation

The trimmed reads were aligned to the assembled transcriptome (length >=200 bp) using Bowtie2 program v2-2.2.6 [71]. We allowed up to 1-mismatches in the seed region (length = 31 bp) and all multiple mapped positions were reported. Of all filtered reads, about ~94% of reads from each sample were properly aligned back to the assembled transcriptome.

### Differential gene expression analysis

Following transcript alignment, differential gene expression analysis was performed using edgeR program [72] for identification of genes that were up-regulated and down-regulated in each crop during tissue culture-based regeneration. Levels of gene expression were represented as log2 ratio of transcript abundance between control and cultured samples. Differentially expressed genes identified in control and cultured samples were analyzed by hierarchical clustering. A heat map was constructed using the log-transformed and normalized value of genes based on Pearson uncentered correlation distance as well as based on complete linkage method.

### Annotation of differentially expressed transcripts

The assembled transcripts were annotated using in-house pipeline (CANoPI – Contig Annotator Pipeline, unpublished) for *de novo* transcriptome assembly. Assembled transcripts were compared with NCBI plant non-redundant protein database using BLASTX program. Matches with an E-value cutoff of 10^−5^ and % identity cutoff of 40% were retained for further annotation. The top significant blast for each of the transcripts was considered for annotation and each of the differentially expressed transcripts were annotated and the organism name was extracted. The predicted proteins from BLASTX were annotated against NCBI plant redundant database, UniProt database and pathway information from other databases like Plant Metabolites Network database. Furthermore, gene ontology (GO) terms for transcripts were extracted wherever possible based on UniProt database. GO terms were mapped to molecular function, biological process and cellular components. Finally, the orthologs gene groups among all upregulated and downregulated DEGs were identified using OrthoMCL Tool [73].

### Quantitative real time-PCR (qRT-PCR)

For validation of RNA-Seq data through qRT-PCR, 72 differentially expressed genes obtained from the three crops were selected and the forward and reverse primer sequences along with the resultant amplicon sizes are presented in Additional file 5: Table S16. The sequences were blasted in NCBI, the consensus sequences for the three crops were derived in Clustal W and primers were designed using Primer3Plus software. For qRT-PCR studies, explants cultured for 7, 14 and 21 days along with control from the three crops were used totaling 12 samples per replicate and each experiment included three biological replicates. The total RNA was isolated, and first strand cDNA was synthesized using one µg of RNA with Super Script III first-strand synthesis kit (Invitrogen, USA) from which one µl of 1:10 diluted cDNA was used as template for qRT-PCR. The qRT-PCR reactions were performed on a Light Cycler 96 System (Roche, USA) using the SYBR premix ExTaq™ II (Takara, Japan) in 96-well optical reaction Roche plates. Each reaction contained 5 μl SYBR Green Master, 0.8 μl template cDNA, 0.4 μl each of the primers (10 μM), and 3.4 μl RNase-free water in a total volume of 10 μl. The qRT-PCR profile was as follows, 95 °C (2 min), 40 cycles of 95 °C (5 s), 60 °C (30 s) with fluorescent signal recording and 72 °C for 30 s. The melting curve was obtained using a high-resolution melting profile performed after the last PCR cycle, 95 °C for 15 s followed by a constant increase in the temperature between 65 °C (15 s) and 95 °C (1 s). The actin gene served as an endogenous control for normalization, and relative fold-changes of the differentially expressed genes were calculated using the ΔΔ cycle threshold (CT) method (2^−^ (ΔC_T_ treatment – ΔC_T_ control) according to Livak and Schmittgen [74].

## Abbreviations

BA: N^6^–Benzyladenine
BP: Biological processes
CC: Cellular components
CPM: Counts per million
2,4-D: 2,4-dichlorophenoxyacetic acid
DEG: Differentially expressed genes/transcripts
FPKM: Fragments per kilobase per million mapped fragments
GO: Gene ontology
IAA: Indoleacetic acid
2-iP: 2-isopentenyladenine
KN: kinetin
MF: Molecular function
MS: Murashige and Skoog
NAA: α-Naphthaleneacetic acid
TDZ: 1-Phenyl-3-(1,2,3-thiadiazol-5-yl) urea (thidiazuron)

## Declarations

### Ethics approval and consent to participate

Not applicable

### Consent for publication

Not applicable

### Availability of data and material

The datasets generated and analysed during the current study are available in NCBI and can be accessed with NCBI accession number PRJNA415556.

### Competing interests

The authors declare that they have no competing interests.

### Funding

The work was carried out at ICAR-IIOR without any funding support.

### Authors’ contributions

SSP was involved in tissue culture work, RNA isolation and manuscript preparation; TM in tissue culture experiments, data recording, RNA isolation, qRT-PCR experiments; PAVT in data interpretation, analysis and draft manuscript preparation; KOA in supervision and guidance, finalization of the technical programme; NC, SG, VKV, KMAVSK, SPL, BK, RVBL in sequencing, transcriptome analysis, bioinformatic analysis, data interpretation and manuscript preparation; SM in conceiving the idea, work plan, interpretation, data analysis and manuscript preparation. All authors read and approved the final manuscript.

## Acknowledgements

SSP, TM and SM thank the Director, ICAR-IIOR for providing the necessary facilities for carrying out the research work.

## Additional files

**Additional file 1: Table S1.** List of all transcripts identified in control and cultured samples of castor with their basic information collected from NCBI, Uniport database along with structural and functional information.

**Additional file 2: Table S2.** List of all transcripts identified after cd-hit-est with 0.9 identity filtering, in control and cultured samples of Jatropha with their basic information collected from NCBI, Uniport database along with structural and functional information.

**Additional file 3: Table S3.** List of all transcripts identified after cd-hit-est with 0.9 identity filtering, in control and cultured samples of sunflower with their basic information collected from NCBI, Uniport database along with structural and functional information.

**Additional file 4: Table S4.** Significant upregulated genes (P ≤ 0.05) identified after DEGs analysis between castor control and cultured genes with their basic information collected from NCBI, Uniport database along with gene ontology and functional role in different metabolic pathways.

**Additional file 4: Table S5.** Significant downregulated genes (P ≤ 0.05) identified after DEGs analysis between castor control and cultured genes with their basic information collected from NCBI, Uniport database along with gene ontology and functional role in different metabolic pathways.

**Additional file 4: Table S6.** Significant upregulated genes (P ≤ 0.05) identified after DEGs analysis between Jatropha control and cultured genes with their basic information collected from NCBI, Uniport database along with gene ontology and functional role in different metabolic pathways.

**Additional file 4: Table S7.** Significant downregulated genes (P ≤ 0.05) identified after DEGs analysis between Jatropha control and cultured genes with their basic information collected from NCBI, Uniport database along with Gene ontology, functional role in different metabolic pathways.

**Additional file 4: Table S8.** Significant upregulated genes (P ≤ 0.05) identified after DEGs analysis between sunflower control and cultured genes with their basic information collected from NCBI, Uniport database along with gene ontology and functional role in different metabolic pathways.

**Additional file 4: Table S9.** Significant downregulated genes (P ≤ 0.05) identified after DEGs analysis between sunflower control and cultured genes with their basic information collected from NCBI, Uniport database along with gene ontology and functional role in different metabolic pathways.

**Additional file 4: Table S10.** List of commonly upregulated groups in castor identified after performing orthologues identification in upregulated DEGs.

**Additional file 4: Table S11.** List of commonly upregulated groups in jatropha identified after performing orthologues identification in upregulated DEGs.

**Additional file 4: Table S12.** List of commonly upregulated groups in sunflower identified after performing orthologues identification in upregulated DEGs.

**Additional file 4: Table S13.** List of commonly downregulated groups in castor identified after performing orthologues identification in downregulated DEGs.

**Additional file 4: Table S14.** List of commonly downregulated groups in jatropha identified after performing orthologues identification in downregulated DEGs.

**Additional file 4: Table S15.** List of commonly downregulated groups in sunflower identified after performing orthologues identification in downregulated DEGs.

**Additional file 5: Table S16.** List of genes validated through qRT-PCR along with the primer sequences and the relative fold change values at four different time points in castor, Jatropha and sunflower.

